# Filamentation and proline inhibition of glutamate kinase

**DOI:** 10.1101/2024.09.19.614007

**Authors:** Tianyi Zhang, Qingqing Leng, Huan-Huan Hu, Ji-Long Liu

## Abstract

Glutamate kinase (GK) is the first committed enzyme in the proline biosynthesis pathway. Belonging to amino acid kinase (AAK) superfamily, most prokaryotic GKs have an additional PseudoUridine synthase and Archaeosine transglycosylase (PUA) domain at the C-terminus, while the function of the PUA domain in GK is poorly understood. Here, we find that *Escherichia coli* GK (EcGK) assembles into filaments and bundles in the state of apo and proline binding. Using cryogenic electron microscopy, we determine the high-resolution structures of EcGK filaments and bundles. The PUA domain is necessary for EcGK filaments and bundles, and the main interfaces have been clearly defined. The feedback inhibitor proline binds at the same pocket as substrate glutamate, inducing conformational changes on nearby regulatory loop which facilitate proline binding. The PUA domain stabilizes the regulatory loop and contributes to proline feedback inhibition. This study reports the special filament-based assembly of EcGK at apo and proline binding state. The first proline binding structure in the GK family illustrates the feedback inhibition mechanism. Intriguingly, the PUA domain is involved in both filamentation and feedback inhibition of EcGK, revealing the versatility of this ancient domain.

## Introduction

Synthesizing proline from glutamate is a conserved anabolic pathway in organisms. In prokaryotes, three enzymes are involved in this pathway: γ-glutamate kinase (GK), γ-glutamyl phosphate reductase (GPR), and pyrroline-5-carboxylate reductase (P5CR). Glutamate will be first phosphorylated to glutamate-5-phosphate (G5P) by GK, and then be reduced to glutamate-5-semialdehyde (GSA) by GPR. GSA will spontaneously cyclize to pyrroline-5-carboxylate (P5C). Finaly, P5C will be converted to proline by P5CR. In many eukaryotes including animals and plants, the first two enzymes GK and GPR merge into a bifunctional enzyme pyrroline-5-carboxylate synthase (P5CS). In this pathway GK catalyzes the first and rate-limiting reaction, which is feedback inhibited by the final product proline[1] **(Figure 1a)**.

**Figure 1.**
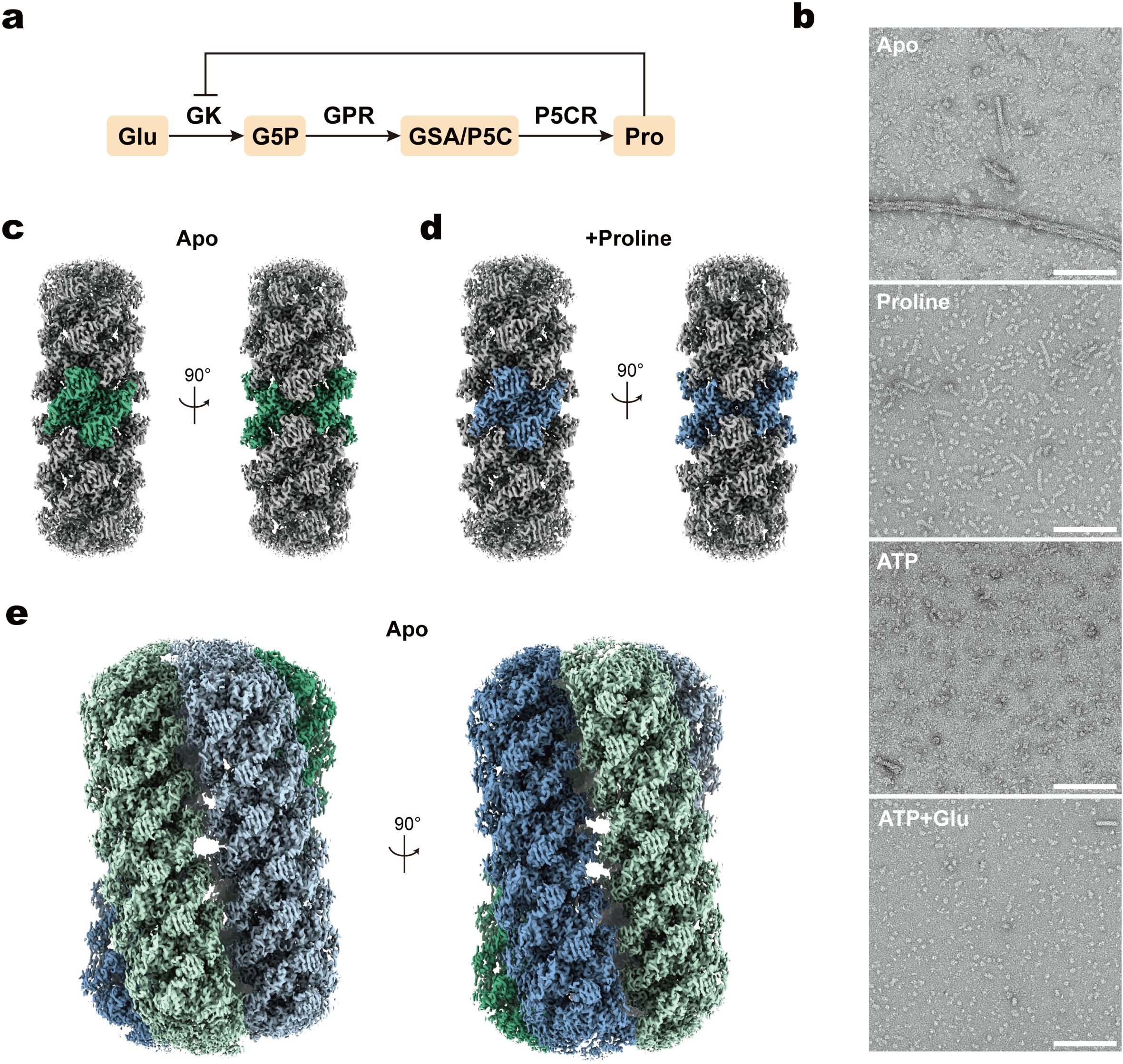
Overview of EcGK function and structure. a. Biosynthesis pathway of proline from glutamate in *E. coli*. Text in orange boxes denote the metabolites in the pathway. Text on arrows indicate the enzymes that catalyze each step of the pathway. “⊥” at the end of line indicates feedback inhibition. b. Negative staining results of EcGK at different conditions. Scale bar: 150 nm. The concentration used: proline 0.5mM, glutamate 20mM, ATP 5mM. c. Cryo-EM density map of the EcGK apo-bundle, composed of four filaments which are indicated with different colors. d. Cryo-EM density map of the EcGK apo-filament. The central tetramer is highlighted with green. e. Cryo-EM density map of the EcGK proline-filament. The central tetramer is highlighted with blue.

Regulation of GK is important for metabolic homeostasis. Hyperactivation of GK kills *Mycobacterium tuberculosis*, thus providing a new strategy for developing antibiotics[2]. Some strains like *Corynebacterium glutamicum* are used for industrial proline production, and removal of feedback inhibition in *Corynebacterium glutamicum* GK (CgGK) improves productivity of proline[3, 4]. Despite the important role of GK feedback inhibition in basic metabolism and industrial application, structural information of proline binding and inhibition mechanism is still limited.

Enzymological properties of wild type and mutant GK gave some insights into the mechanisms of proline inhibition. The concentration of glutamate required for 50% activity of GK increases when increasing proline concentration, indicating proline is a competitive inhibitor for glutamate[5]. Molecular docking results suggest a similar binding mode for proline and glutamate and consist with the effect of D148 to D150 mutants[6]. However, many mutations which decrease proline inhibition cluster in the loop E135-D148, and some of these mutations are useful for industrial applications[1, 4, 7]. Effects of these mutations on proline inhibition cannot be well illustrated with current crystal and molecular docking models.

Some interesting properties of *Escherichia coli* GK (EcGK) have been found during the investigation of proline inhibition. When applied to size exclusion chromatography (SEC), proline binding EcGK shows increased molecular weight compared with APO EcGK, indicating EcGK may aggregate to some kinds of high order structure[8]. Recent studies on P5CS give some clues on this aggregation. In *Drosophila*, P5CS forms cytoophidia *in vivo* and metabolic filaments *in vitro*[9–11]. For animal P5CS, two interfaces on GK and GPR domains contribute to the assemble of filament[10]. For plant P5CS, the stable helical interface only locates on GK domain, and GK domain itself can form filament[12]. Both animal and plant P5CS forms filaments which promote their activities. Given the homology of prokaryotes GK and GK domain of P5CS, it is possible that EcGK also form filament.

Many enzymes have been reported to form evolutionary conserved filament[13–15]. For example, structures of CTPS and PRPS filaments have been characterized exhaustively, and their filament interfaces are conserved across a wide range of organisms[16–23]. But for prokaryotes GK and eukaryotes P5CS, some significant changes occurred during evolution. Most GKs have an additional PseudoUridine synthase and Archaeosine transglycosylase (PUA) domain[24, 25] at C-terminus, while PUA domain is completely lost in P5CS[26]. PUA domain is an ancient RNA binding domain, but no evidence shows the GK could form complexes with RNA[25, 27]. PUA domain is not necessary for activity of GK, and its functions in GK remain largely uncertain[8]. In P5CS, a motif named “hook” located in GK domain is important for filamentation, but in EcGK, the corresponding region is significantly shorter[10, 12]. If EcGK also forms filament, the assembly and interfaces for EcGK filament should be different from *Drosophila melanogaster* P5CS (DmP5CS) and *Arabidopsis thaliana* P5CS (AtP5CS). These differences make GK and P5CS an interesting model for studying the evolution of metabolic filaments.

In this work, we find that purified EcGK also forms filaments and bundles. Unlike P5CS filaments that promote enzyme activity, EcGK filaments disassemble during the catalytic process. We have solved the high-resolution structure of EcGK apo-bundle, apo-filament, and proline-filament. The filament interfaces of apo-filament and proline-filament are similar, but completely different from those of P5CS. Compared with previously published EcGK models, the PUA domain rotates around amino acid kinase (AAK) domain, forming new filament and bundle interfaces. Surprisingly, the interaction between the PUA domain and the AAK domain promotes proline feedback inhibition. By resolving the first proline binding GK structure, we reveal the mechanism of effective inhibition of proline. Our results indicate that EcGK utilizes the PUA domain for metabolic regulation, revealing the versatility of this ancient domain.

## Results

### EcGK forms filaments and bundles under specific conditions

We used the T7-based overexpression system for producing EcGK **(Supplementary Figure 1 a)**. N-terminal 6×his-SUMO tag was used for purification. After SUMO protease digestion, we acquired highly purified EcGK without excessive residues (**Supplementary Figure 1 b, c**). To investigate polymerization state of EcGK, we made negative staining samples of EcGK after incubated at different conditions **(Figure 1 b)**. At apo state (no ligand binding state) EcGK forms filaments and bundles of filament. Proline impedes the formation of bundles but allows the formation of filaments. At low ATP concentrations (1mM, 2mM), the formation of filaments and bundles did not seem to be affected (**Supplementary Figure 2**). However, higher ATP concentration (5mM) will disrupt almost all filaments and bundles **(Figure 1 b)**. But the addition of proline can rescue filaments disassembly (**Supplementary Figure 2**). The filaments and bundles still exist after incubation with glutamate (**Supplementary Figure 2**). Although filaments almost completely disappeared after incubation with all substrates (ATP, Glutamate) **(Figure 1 b)**, the addition of proline was able to restore filament formation (**Supplementary Figure 2**). The existence of proline can sustain the formation of GK filaments.

To investigate the structural basis of EcGK filaments and bundles, we prepared cryogenic EcGK samples at apo state and incubated with proline. Using cryo-EM, we resolved the structure of EcGK apo-filament, apo-bundle and proline binding filament to 2.67 Å, 2.97 Å and 2.76 Å, respectively (**Supplementary Table 1, Supplementary Figure 3-5**). The apo-filament and proline binding filament have similar conformations, and both filaments are assembled by stacking of tetramers (**Figure 1 c-d**). The apo-bundle is composed of four slightly bent filaments (**Figure 1 e**).

### The structure of EcGK apo-filament reveals novel filament interfaces

When forming filament, the relative position of AAK domain and PUA domain is different from previously published EcGK tetramer models (**Figure 2 a**). In published models like 2J5T, the PUA domain and AAK domain locate on a same plane, forming a flat conformation (**Supplementary Figure 6 a**). While in the EcGK filament, PUA domain rotates away from this plane and the tetramer is rearranged to a crossed conformation (**Supplementary Figure 6 a**). In both models, the AAK domain, linker and PUA domain interact with each other to stabilize the conformation. For the crossed conformation, the main interactions are π-π/cation-π interactions among W268, R111, F114, R118 and hydrogen bonding/electrostatic interactions among R118, D363, N116, D119, R309, R329 (**Supplementary Figure 6 b**). In the flat conformation, the W268 mediated π-π/cation-π interactions are weakened, and the potential interactions only occur between W268 side chain and backbone/side chain of R111 (**Supplementary Figure 6 c**). A different set of hydrogen bonding/electrostatic interactions among D222, R367, R226, K266, R361, M364 stabilize the flat conformation (**Supplementary Figure 6 c**).

**Figure 2.**
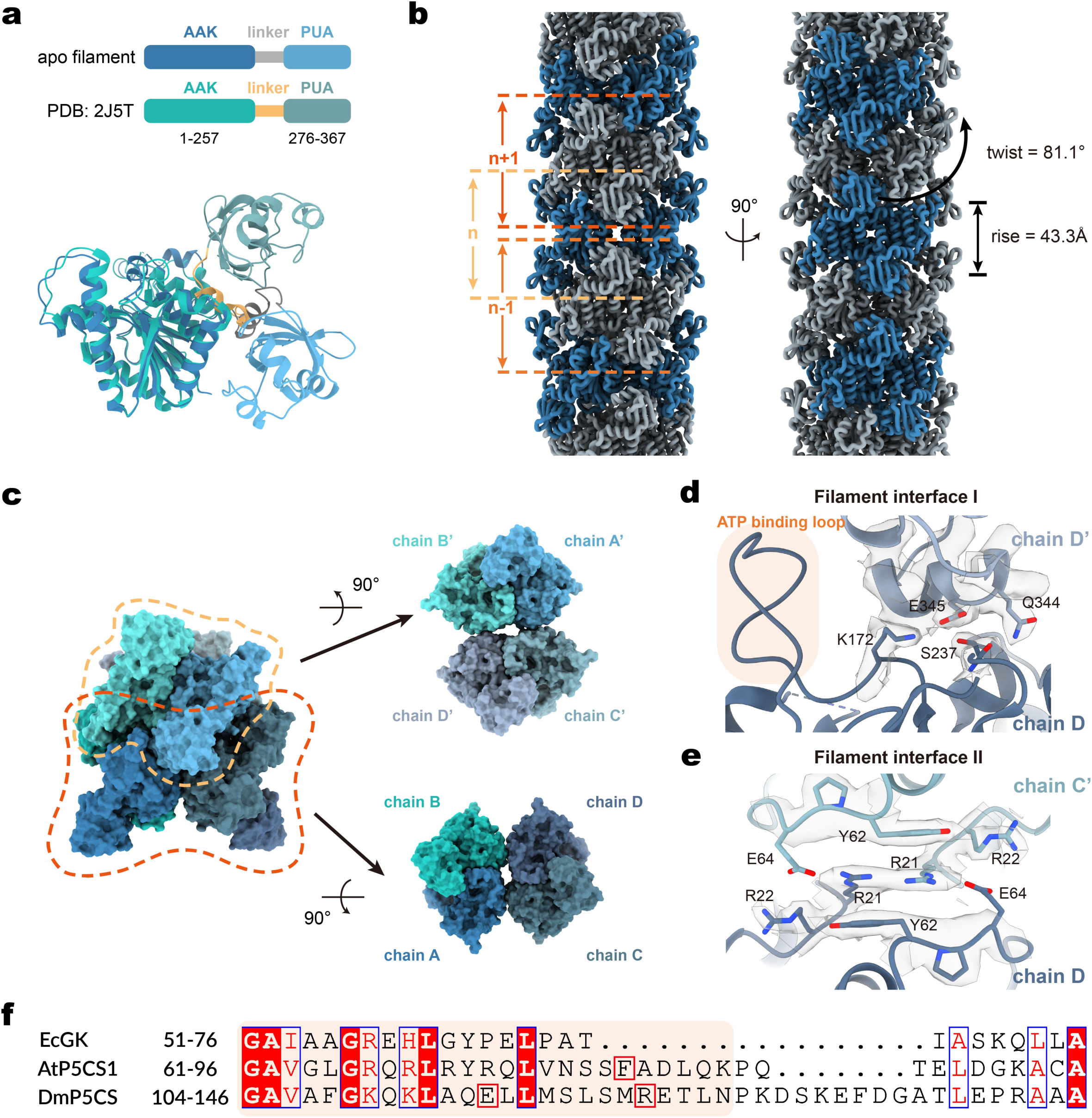
Structural analysis of EcGK apo-filament. a. Comparison of monomer from EcGK apo-filament and published model 2J5T. AAK domain of 2J5T monomer is aligned and superimposed onto apo-filament monomer. The AAK domain, linker, and PUA domain are colored as the scheme shown on the top. The residue ranges of AAK domain and PUA domain are labeled below the color scheme. b. Architecture of EcGK apo-filament. Tetramers are colored alternately in blue and gray. c. Octamer from EcGK apo-filament. Two tetramers are separated and rotated to show the interfaces. d. Zoom-in view of interface I. Density map around interested residues are shown as transparent grey. ATP binding loop is labeled with light orange background. e. Zoom-in view of interface II. Density map around interested residues are shown as transparent grey. f. Part of sequence alignment results of EcGK, AtP5CS1, and DmP5CS. Residues corresponding to hook are labeled with light orange background. F80 in AtP5CS1 and R124 in DmP5CS are labeled with red boxes.

The basic helical unit of EcGK apo-filament is tetramer with D2 symmetry. Many tetramers densely stack along the axis which passes through center of tetramer with 43.3 Å rise and 81.1° twist (**Figure 2 b**). By applying helical symmetry to a tetramer composed of chain A, B, C, and D for one time, we get a copy of this tetramer composed of chain A’, B’, C’, and D’ that are corresponding to original chain A-D. These two tetramers assemble to an octamer which can be used to illustrate all interfaces between helical units (**Figure 2 c**). Two main helical interfaces are found in apo-filament. The interface I lies between chain X and X’ (X = A, B, C, D) and connect PUA domain from one chain and AAK domain from another chain. This interface locates near the ATP binding loop which recognizes the adenine of ATP (**Figure 2 d**). Salt bridge between K172 and E345 and hydrogen bond between S237 and Q344 are main interactions (**Figure 2 d, Supplementary figure 6 d**). Other weak interactions may also contribute to this interface, like π-π interactions between R180 guanidine group and L350-G351 peptide amide group, R337 guanidine group and P179-R180 peptide amide group, and cation-π interactions between R337 and Y175 (distance between R337 NH and Y175 CZ is 5.2 Å) (**Supplementary figure 6 e**). The interface II is a sandwich structure and lies between AAK domains of chain A and chain B’ (and chain D and chain C’) (**Figure 2 e**). The main interactions are π-π/cation-π interactions network among R21 and Y62 (**Supplementary figure 6 f**). Salt bridge between R21 and E64 may also stabilize this interface (**Supplementary figure 6 f**). However, the density of E64 side chain is completely lost in this map, making the position of E64 side chain and this salt bridge interaction uncertain (**Figure 2 e**). E64 may also contribute to this interface through electrostatic interaction with R22 (**Figure 2 e**).

The interfaces of EcGK filament are quite different from those in P5CS. A motif named “hook” forms the helical interfaces in GK domain of P5CS. In AtP5CS1 and AtP5CS2, F80 on the hook is the key residue for this interface. In DmP5CS, E116 and R124 on the hook interact with each other to stabilize the filament. The hook of EcGK is much shorter than AtP5CS and DmP5CS (**Figure 2 f**). And residues in EcGK corresponding to AtP5CS F80 or DmP5CS E116 and R124 are not conserved (**Figure 2 f**). In both AtP5CS and DmP5CS, four hooks form one interface. While in EcGK, four hooks and four L60-T68 loops form two same interfaces. In addition, PUA domain also contribute to the helical interface of EcGK, and this domain is lost in P5CS.

### The structure of EcGK apo-bundle

Using the same EcGK apo dataset, we also reconstructed apo-bundle (**Supplementary Figure 4**). Intuitively, EcGK apo-bundle consists of four apo-filaments (**Figure 1 e**). All four filaments are identical and slightly bent to be twisted together like a rope (**Figure 3 a, b**). The helical symmetry of apo-filament is broken because of the curvature. Filament in apo-bundle is left-hand helix in which the rise is 84.0 Å and twist is –16.7° (**Figure 3 a, b**). Furthermore, the whole bundle can be described by octamer and one-set of helical symmetry, get rid of the concept of filament (**Figure 3 a, c**). The bundle is a right-hand helix, and the rise and twist are 21.0 Å and 85.9° respectively (**Figure 3 a**). After four turns, the fifth octamer contacts with the first octamer and they belong to the same filament.

**Figure 3.**
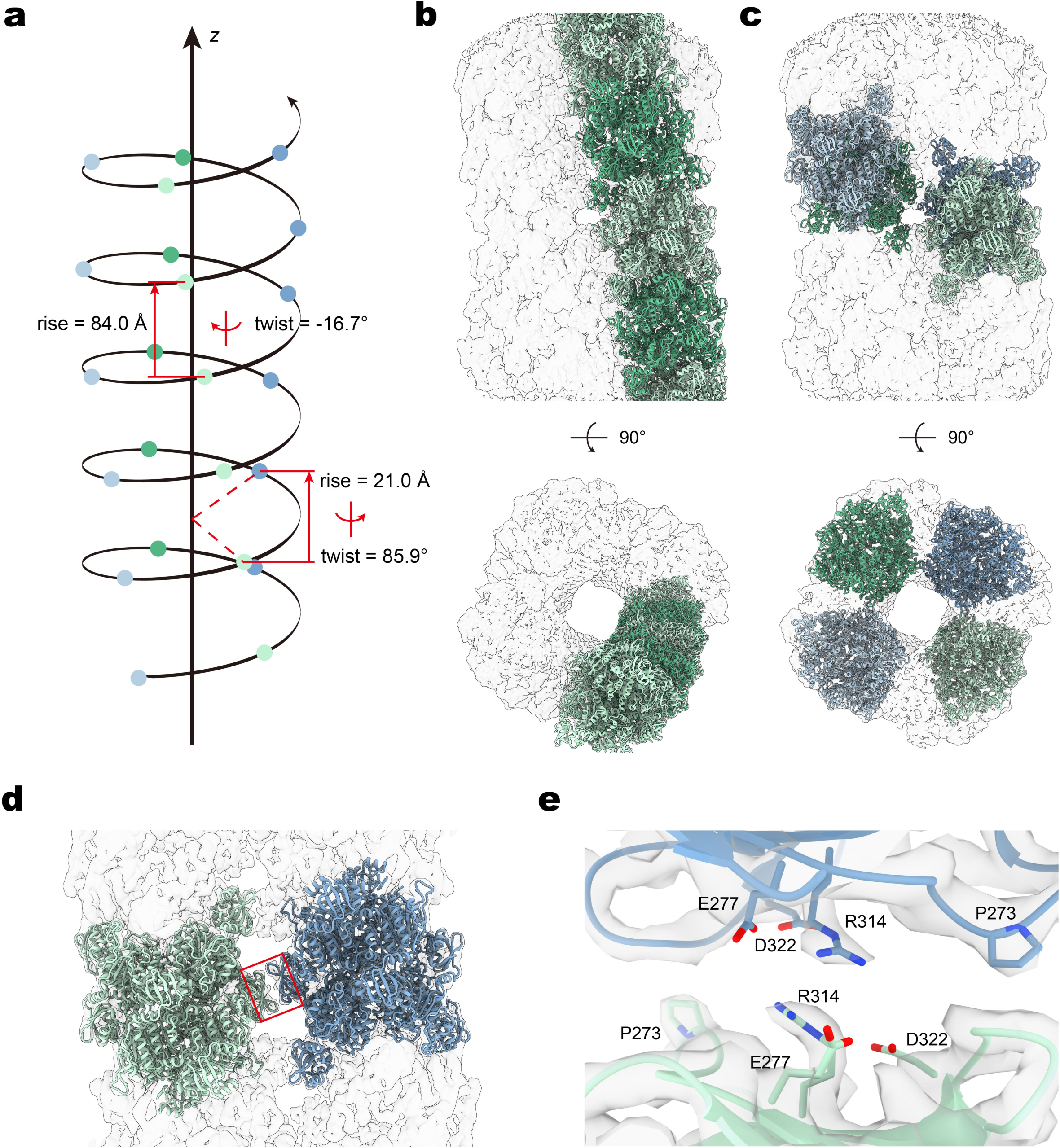
Structural analysis of EcGK apo-bundle. a. Helical symmetry display of EcGK apo-bundle. Each circle represents an octamer. Octamers from the same filament are indicated with the same color. b. EcGK apo-bundle map with one filament model. c. EcGK apo-bundle map with four octamer models. d. Tow octamer models fitted in EcGK apo-bundle map. Bundle interface is indicated with red box. e. Zoom-in view of EcGK apo-bundle interface.

There is only one kind of interface between filaments. This bundle interface is mediated by R314, E277 and D322 on the PUA domain (**Figure 3 d, e**). The density of R314 side chain is well resolved but density for E277 and D322 carboxyl groups is largely lost (**Figure 3 d**). The guanidine group of two R314 residues are perfectly parallel and the distance between two CZ atoms sustain the restriction of π-π interaction (**Supplementary figure 7 a**). E277 and D322 may contribute to this interface either by neutralizing the charge of R314 and reducing the repulsion between two R314 through intramolecular electrostatic interactions, or by directly attracting R314 through intermolecular electrostatic interactions. Due to the lack of density the distance between E277/D322 carboxyl groups and R314 guanidine group cannot be measured accurately, but in current model both intramolecular and intermolecular electrostatic interactions are feasible. For intramolecular interactions, the distances between D322 OD atoms and R314 NE atom fall in the range of 3 Å to 4 Å, and the distances between E277 OE atoms and R314 NE/NH atoms fall in the range of 4 Å to 5 Å (**Supplementary figure 7 b**). For intermolecular interactions, the distances between E277/D322 carboxyl O atoms and R314 NH atoms fall in the range of 4 Å to 5 Å (**Supplementary figure 7 c**). R314A mutation abolishes the formation of EcGK bundle, confirming the importance of R314 on bundle interface (**Supplementary figure 7 d, e**).

### Proline binding induces conformational changes in filaments

After incubated with 0.5 mM proline, we made proline state sample and reconstructed EcGK proline-filament (**Supplementary Figure 5**). In the same dataset no EcGK bundle was found (**Supplementary Figure 8**). For the first time we observed the binding mode of proline in GK. The imino group of proline forms salt bridge with D137, and the carboxyl group forms four hydrogen bonds with G51, A52, I53 and N134 (**Figure 4 a**). Proline binding induces conformational changes compared with apo-filament. There are two significant changes in AAK domain: the E135-N149 loop and the I69-Y94 helix accompanied with adjacent L60-T68 loop (**Figure 4 b**). Different with previously published glutamate binding structure, a short helix (A142-V146) forms within the E135-N149 loop in both apo-filament and proline filament. At apo state, K145 forms salt bridge with D137 (**Supplementary Figure 9 a**). After proline binding, this short helix flips and enhances interactions within a dimer. Referring two chains in a dimer as chain A and chain B, K145 from chain A interacts with PUA domain from chain B through cation-π interaction with Y352 and hydrogen bond with Y354 (**Figure 4 c**). The I69-Y94 helix is more curved after proline binding, and the adjacent L60-T68 loop shifts towards the same direction. New hydrogen bond forms between E353 and I69 backbones, and Q73 shifts the hydrogen bond from A140 to D137 (**Figure 4 c, Supplementary Figure 9 a**). Taken together, after proline binding new dimer interface is formed between PUA domain and AAK domain. Two PUA domain in one dimer rotate toward each to form a tighter dimer (**Figure 4 d**).

**Figure 4.**
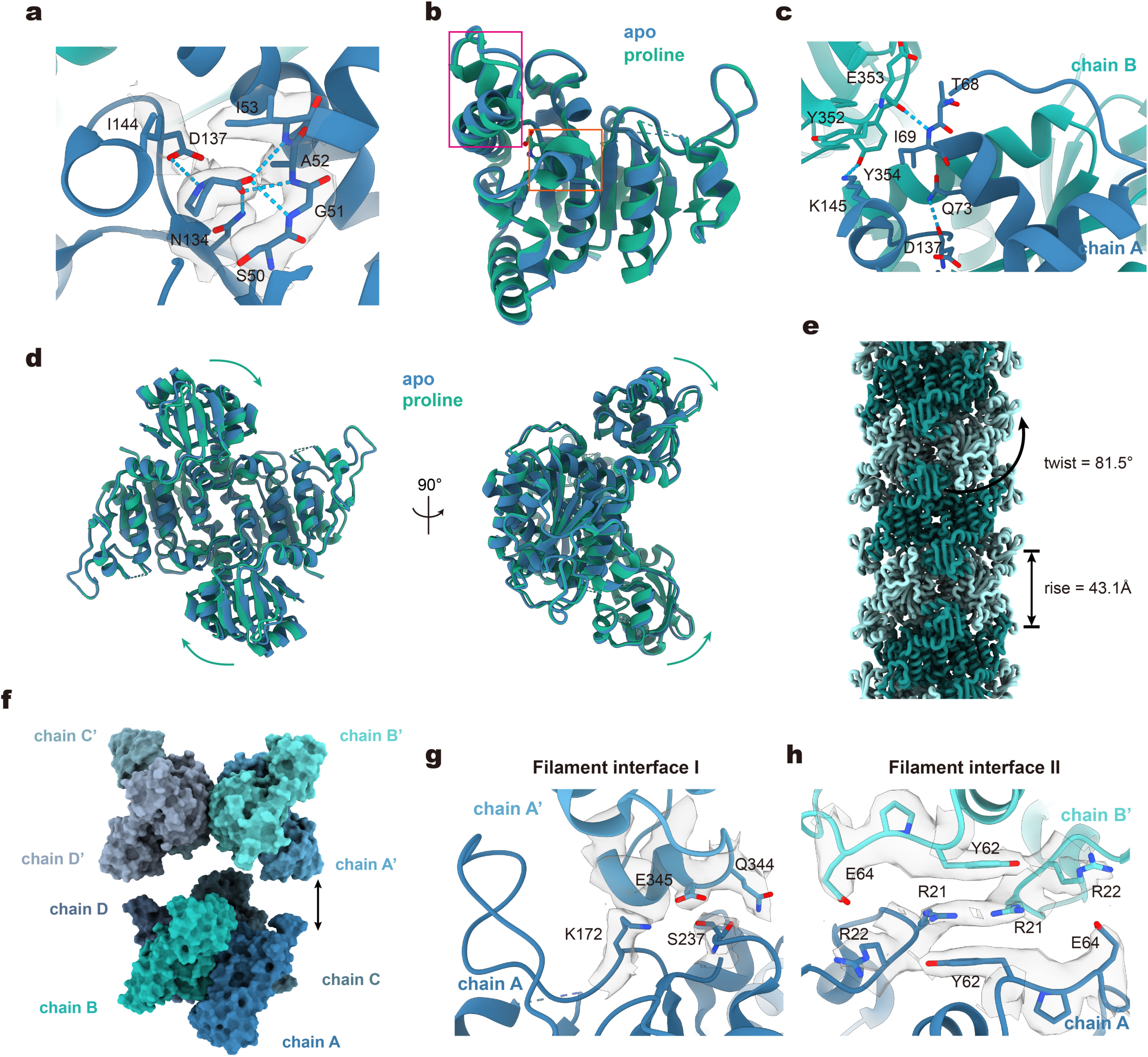
Structural analysis of EcGK proline-filament. a. Detailed interactions analysis of proline binding. Density for proline and nearby residues are shown as transparent grey. Hydrogen bonds/salt bridges are indicated by blue dashed lines. b. Superposition of monomers from apo-filament and proline-filament. Magenta box and orange box indicate the differences between two models. c. Details of the PUA-AAK interface within one dimer. Hydrogen bonds/salt bridges are indicated by blue dashed lines. d. Superposition of dimers from apo-filament and proline-filament. Rotation of proline-filament relative to apo-filament are indicated by green arrows. e. Architecture of EcGK proline-filament. Tetramers are colored alternately in dark green and light green. f. Octamer from EcGK apo-filament. Two tetramers are separated to show the interfaces. g-h: Zoom-in view of g) interface I; and h) interface II. Density map around interested residues are shown as transparent grey.

The overall structure of EcGK proline-filament is quite similar to apo-filament except the filament interface II. The twist and rise are slightly changed upon proline binding (**Figure 4 e**). Like apo-filament, all filament interfaces in proline-filament can be well illustrated through an octamer (**Figure 4 f**). Filament interface I remains unchanged comparing to apo-filament (**Figure 4 g**). However, filament interface II becomes different due to the shift of L60-T68 loop. Two Y62 get closer to each other, therefore the intermolecular interactions (for example, chain A and chain B’) between Y62 and R21 are enhanced (**Figure 4 h, Supplementary Figure 9 b**). The density of E64 side chain is clear in proline-filament, and based on the distances E64 has electrostatic interaction with R22 but probably has no direct interaction with R21 (**Figure 4 h, Supplementary Figure 9 c**).

### The PUA domain promotes proline feedback inhibition

In agreement with previous speculation, the binding site of proline and glutamate is largely overlapped (**Figure 5 a**). The binding modes of α-carboxyl groups and amino (imino) groups in glutamate and proline are almost identical, while the different side chains induce two conformations for the E135-N149 loop. Proline is a compact amino acid, and it stabilizes the E135-N149 loop at a “close” conformation. The pocket for proline is much deeper and tighter than the pocket for glutamate (**Figure 5 b, c**). The A142-V146 short helix covers the top of proline, and on the other side, D148 forms hydrogen bonds with T13 and S50 to shield proline, leaving only a small entrance for this proline pocket (**Figure 5 b, d**). In glutamate binding structure the E135-N149 loop is stabilized at a “open” conformation and no short helix is formed. D148 rotates outwards the pocket since the γ-carboxyl group of glutamate occupies the position.

**Figure 5.**
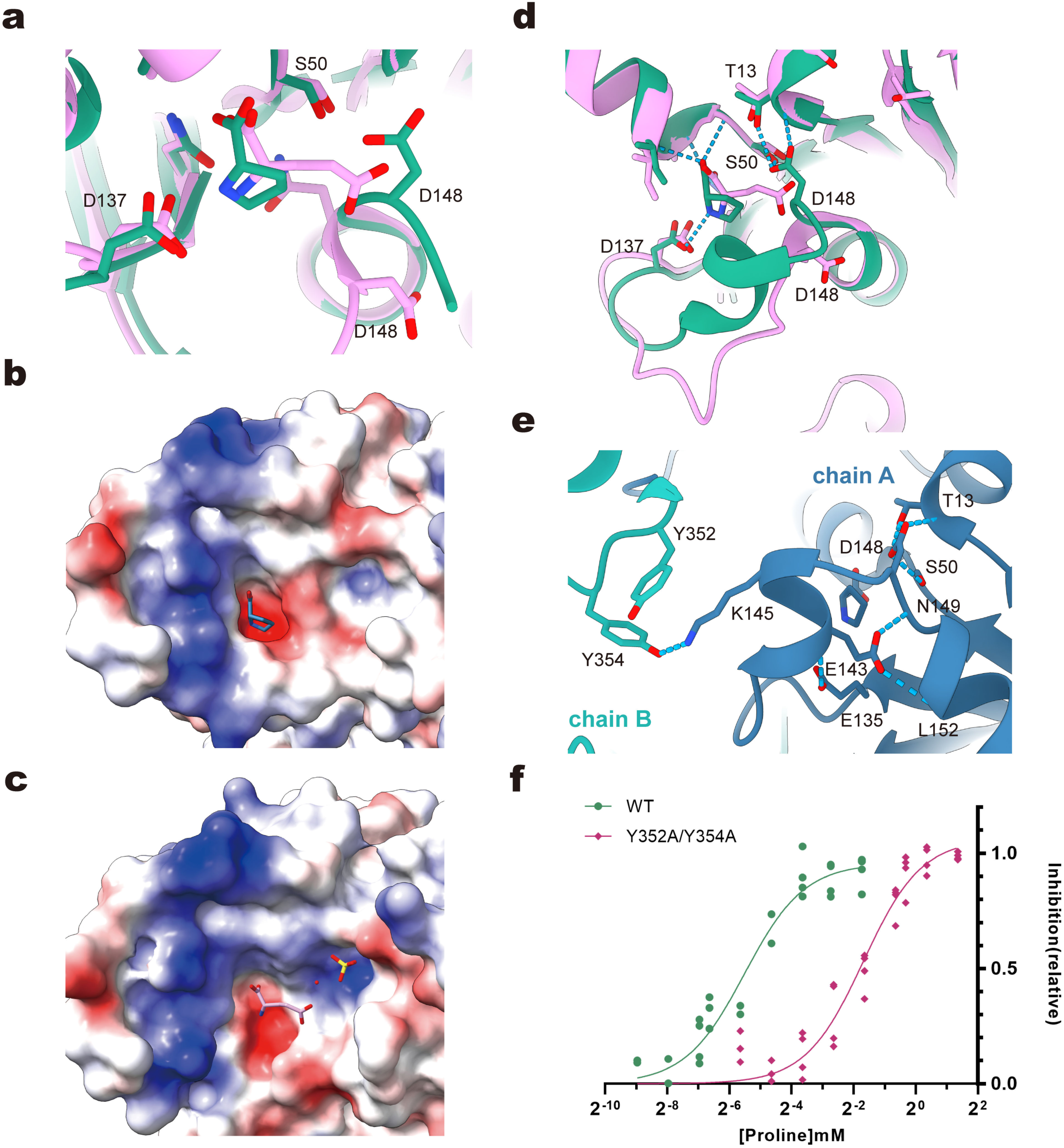
a. Structure comparison of proline and glutamate binding pocket. The structure of proline binding is colored in green and the glutamate binding structure (PDB:2J5T) is color in pink. b. Electrostatic potential surface of proline binding pocket. c. Electrostatic potential surface of glutamate binding pocket. d. Detailed interactions analysis near the proline binding pocket. Colored as the same scheme in panel a. e. Detailed interactions analysis of E135-N149 loop and PUA domain. Chain A and chain B form a dimer. Chain A is colored in deep blue, the PUA domain of chain B is colored in cyan. f. Effects of EcGK-Y352A/Y354A mutant on the concentration requirement of proline feedback inhibition. Values are expressed relative to the value of complete inhibition.

The close conformation for E135-N149 loop stays in a low energy state. Besides the backbone hydrogen bonds in the short helix, side chains of E135, E143 and E148 also form hydrogen bonds to stabilize this conformation. Interestingly, K145 in the short helix could interact with PUA domain from another chain in a dimer through Y352 and Y354 (**Figure 5 e**). To test if these interactions facilitate proline inhibition, we constructed the EcGK-Y352A/Y354A mutant. As expected, the requirement of proline for 50% inhibition increased about 15-fold for this mutant (**Figure 5 f**). Taken together, proline binding locks EcGK in a low energy state, building a high energy barrier to prevent the competitive binding of glutamate. This explains the sensitivity of proline feedback inhibition for EcGK. PUA domain helps stabilize the close conformation of E135-N149 loop and further promotes proline feedback inhibition.

### The PUA domain is necessary for filamentation

PUA domain provides interfaces for both filament and bundle. PUA domain is necessary for the assembly of bundle from filament, which has demonstrated by EcGK-R314A mutant. However, two interfaces contribute to filament assembly, and it is unclear if AAK domain itself is enough for filamentation. To explore this possibility, we constructed PUA domain deletion mutant named “truncated-EcGK” in which EcGK is truncated after residue 274. During purification, truncated-EcGK shows a sharp UV280 absorption peak and smaller molecular weight in size exclusion chromatography (**Supplementary Figure 10**). The negative staining results show that the truncated mutant cannot form filaments or bundles anymore (**Figure 6 a**). Thus, PUA domain is necessary for filaments and bundles formation.

**Figure 6.**
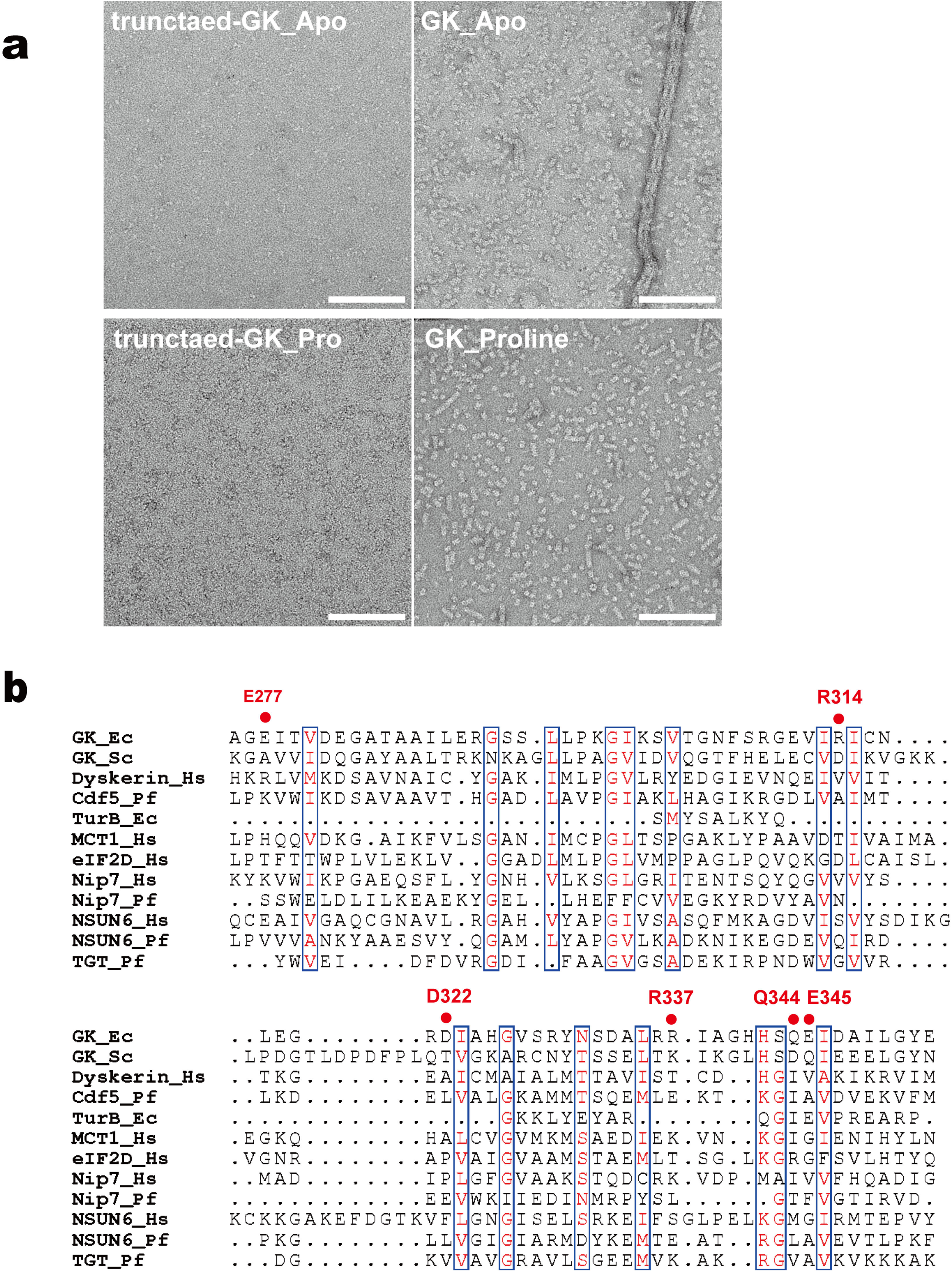
a. Negative staining results of EcGK at different conditions. Scale bar: 150 nm. The concentration used: proline 0.5mM. b. Sequence alignment of 12 proteins contain PUA domain. Ec, E. coli; Sc, S. cerevisiae; Hs, H. sapiens; Pf, P. furiosus.

In previous studies PUA domain is known as an RNA binding domain. It is curious if the key residues for EcGK filament and bundle formation are conserved in other PUA containing proteins. We selected 12 proteins containing PUA domain from archaeal, bacterial and eukaryotic, and extracted PUA domains for sequence alignment (**Figure 6 b**). According to structural information we focused on the key residues Q344, E345, R337 in filament interface and E277, D322, R314 in bundle interface. None of these residues is conserved, indicating EcGK utilize PUA domain in a unique way.

## Discussion

In this work, we determine the filament and bundle structures of EcGK in apo and proline binding state, adding new members to the metabolic filament family. Despite reports that the homologous protein P5CS forms cytoophidia *in vivo* and filaments *in vitro*, EcGK filaments are still surprising[9, 10, 12]. Firstly, EcGK forms inactivate filaments at apo or inhibition state, while P5CS forms active filaments that promote P5CS activity. Secondly, the filament interfaces are quite different in EcGK and P5CS. The formation of EcGK filaments does not rely on the slender hook structure in animal or plant P5CS, and on the other hand, tetramers stack through a lager interface between AAK and PUA domains.

The first proline binding structure for GK enables us to analyze the mechanisms for the highly efficient feedback inhibition. EcGK undergoes conformational changes in response to proline binding, leading to tighter binding pocket and lower energy state. So that the competitive binding of glutamate needs to overcome a huge energy barrier. EcGK also forms more compact filament after proline binding due to the rotation of PUA domain and results in a stronger filament interface. Since PUA domain mediates the filamentation of EcGK, the rotation of PUA domain may be coordinated in one filament, and we speculate that after partially proline binding the coordinated filament may facilitate the subsequent binding of proline. This filament structure may help *E. coli* quickly responds to proline feedback inhibition in relative low concentration. This synergistic model for filamentation and feedback inhibition of EcGK is appealing, and further research is needed to explore the relationship between proline feedback inhibition and filaments.

Relying on the elucidation of the proline-bound GK filament structure, we highlight the importance of E135-N149 loop in proline binding and feedback inhibition. It is a good approach to disrupt the proline inhibition on GK by regulating this loop. In fact, there are many applications of GK in industrial proline production, mostly by using some random mutants such as G149, V150 to release the feedback inhibition of proline in CgGK[3, 4]. These mutation sites correspond to E143, K145 in EcGK, which are located on this regulatory loop and destroy the key interactions. In addition, some recent studies found GK can be a new drug target for anti-TB drugs discovery, the potential binding interaction of compounds and Mtb GK was speculated at position R153, F154, corresponding to the residues K145, V146 of EcGK, also on the E135-N149 loop[2]. The specific mechanism of these applications was not clear before, our elucidation of exact regulatory mechanism of proline feedback inhibition on GK supplemented this understanding. This work may provide new insights and solutions for more targeted drugs development and more efficient engineered stains construction.

Finally, it is interesting that the PUA domain modulates both oligomerization and activity of EcGK. These are unconventional functions for this ancient domain, and accordingly, key residues at the filament and bundle interfaces are not conserved in other PUA containing proteins. Although the PUA domain plays important roles in EcGK and most GKs have PUA domains, we should note that approximately 20% of GKs lack PUA domains [26]. The regulation of GK by other species and the gain or loss of PUA domains during evolution remain a mystery for future research.

## Methods

### EcGK plasmid construction and protein purification

The full-length EcGK gene was cloned into a pET28a vector, N-terminal 6×His tag and a following SUMO-tag was fused at the N terminus of EcGK.

The expression vector of EcGK was transformed into E. coli strain Rosetta (DE3). For protein expression, cells were first cultured at 37 ℃ to OD600 = 0.8-1.0, then protein expression was induced with 100 μM IPTG at 16 ℃ overnight (16-18h). Cells were harvested by centrifugation at 4 ℃ and 4000 g, resuspended with lysis buffer (50 mM Tris-HCl pH 8.0, 500 mM NaCl, 10% glycerol, 20 mM imidazole, 1 mM PMSF, 5 mM β-mercaptoethanol, 5 mM benzamidine, 2 μg/ml leupeptin, and 2 μg/ml pepstatin). Then the cell suspension was homogenized with ultrasonic homogenizer (Scientz-IID) or high-pressure homogenizer (Union-Biotech LH-03). The lysate was centrifugated at 4 ℃, 18000 g for 45 min. Supernatant from 1L E. coli medium was incubated with 1 mL Ni-NTA agarose resin (QIAGEN) for 1 h. Then the resin was washed with 8 mL wash buffer containing 40 mM imidazole (with other components same as lysis buffer). Then proteins were eluted with 4-6 mL elution buffer (50 mM Tris-HCl pH 8.0, 500 mM NaCl, 250 mM imidazole, 5 mM β-mercaptoethanol). Yeast ULP1 was used to remove SUMO-tag. About 6 mL of storage buffer (25 mM Tris-HCl pH 8.0 and 150 mM NaCl) was added together with ULP1 for optimal ULP1 activity. After incubated and concentrated at 4 ℃, proteins were further purified through Superose 6 Increase 10/300 GL (Cytiva) in storage buffer (25 mM Tris-HCl pH 8.0 and 150 mM NaCl). Peak fractions were collected, concentrated, dispensed, snap-frozen with liquid nitrogen and stored at –80°C before use.

### EcGK activity Pi assay

The product G5P synthesized by GK can cyclize spontaneously and form phosphate. The reaction buffer contains 25 mM Tris-HCl pH 8.0, 150 mM NaCl, 20 mM monosodium glutamate, 10 mM MgCl2, 5 mM ATP (Takara, sodium salt, pH 7.0). 500nM enzyme was added to start reaction. The reaction system was incubated at 37 ℃. Reaction was terminated at different times by adding ice-cold 4M perchloric acid to final concentration of 1 M. Then perchloric acid was neutralized and precipitated with 2M ice-cold KOH (final molar ratio perchloric acid: KOH = 1: 1). The reaction mixture was centrifugated at 4 ℃, 13000 g for 15 min. Supernatant was used to determine Pi concentration with Malachite Green Phosphate Detection Kit (Beyotime). The A630 is recorded with SpectraMax i3 plate reader (Molecular Devices).

### Negative staining

Proteins were incubated with different substrates on ice before negative staining. All substrates used here had same concentration as in activity assay. Proteins were diluted to about 2μM. After incubation, protein samples were applied to plasma cleaned carbon-coated EM grids (400 mech, EMCN). The grids were then washed in ddH2O for twice and stained with 2% uranyl formate. Negative-stain EM grids were imaged with a Talos L120C microscope (FEI).

### Cryo-EM grid preparation and data collection

For cryo-EM, purified EcGK was diluted to approximately 20 μM in reaction buffer. EcGK sample was incubated in reaction buffer for about 10 min on ice before vitrification. The concentration of proline used in EcGK-proline sample was 0.5mM. The sample was applied on H2/O2 plasma cleaned ANTcryo™ holy support film (Au300-R1.2/1.3) and then was immediately blotted for 3.0 s. Then the grid was plunge-frozen in liquid ethane cooled by liquid nitrogen using Vitrobot (Thermo Fisher) at 8°C with 100% humidity.

Images were collected on Titan Krios G3 (FEI) equipped with a K3 Summit direct electron detector (Gatan), operating in counting super-resolution mode at 300 kV. Automatic data collection was facilitated with SerialEM[28]. For EcGK apo dataset, we used total dose of 60 e−/Å2 subdivided into 40 frames in 2.4 s exposure. For EcGK proline dataset, we used total dose of 50 e−/Å2 subdivided into 35 frames in 2.1 s exposure. The images were recorded at a nominal magnification of 22,500 × and a calibrated pixel size of 1.06 Å, with defocus ranging from 0.8 to 2.5 μm.

### Data processing

All cryo-EM data processing and reconstruction were performed with CryoSPARC v4.5.1[29]. Reconstruction procedures for three structures are similar and details are shown in **Supplementary Figure 3-5**. Generally, we used patch motion correction and patch CTF estimation for pre-processing. Hundreds of particles were picked manually to generate the initial 2D templates. Using a subset of micrographs, we made the first 3D reconstruction and generated high quality 2D templates with this relative high-resolution model. Due to the intrinsic heterogeneity in EcGK apo samples, we cleaned the particles strictly. Besides multiple rounds of 2D classification, we used heterogeneous refinement with given initial volumes to separate filament and bundle, complete particles and broken particles. Some junk particles were removed according to the tilt angles during helical refinement, and the script for this purpose was released on https://github.com/ScientificWitchery/cryoEM_dataprocessing.

### Model building

The published model PDB:2J5T was used as initial model. AAK domain and PUA domain of initial model were separated and fitted in density map using UCSF ChimeraX[30], and the linker was rebuilt manually according to the density using Coot[31]. Models were iteratively refined using a combination of Coot[31] and phenix.real_space_refine in Phenix[32, 33]. UCSF ChimeraX[30] was also used for visualization purpose.

### Data availability

The structure data accession codes are EMD-60346, EMD-60352, EMD-60397, PDB-8ZPJ, PDB-8ZPR and PDB-8ZRI.

## Acknowledgments

EM data were collected at the ShanghaiTech Cryo-EM Imaging Facility. We thank the Molecular and Cell Biology Core Facility at the School of Life Science and Technology, ShanghaiTech University, and Shanghai Frontiers Science Center for Biomacromolecules and Precision Medicine for providing technical support. This work was supported by grants to J.-L.L. from the Ministry of Science and Technology of China (number 2021YFA0804700), National Natural Science Foundation of China (number 31771490), Shanghai Science and Technology Commission (number 20JC1410500), UK Medical Research Council (grant numbers MC_UU_12021/3 and MC_U137788471).

## Author Contributions

J.L.L. and T.Z. designed the research; H.H.H. reconstructed the initial expression system and tested the protein purification; T.Z. and Q.L. improved the expression system, purified protein, prepared EM samples, and collected EM data; T.Z. processed the cryo-EM data. Q.L. performed the enzyme assays. T.Z. and Q.L. analyzed the data and wrote the manuscript; J.L.L revised manuscript, obtained the funding and supervised the project.

## Competing Interests Statement

Authors declare that they have no competing interests.

---

**Supplementary Table 1.** Cryo-EM data collection and model refinement.

|  | EcGK apo-bundle<br>(EMD-60346, PDB 8ZPJ) | EcGK proline-filament<br>(EMD-60352, PDB 8ZPR) | EcGK apo-filament<br>(EMD-60397, PDB 8ZRI) |
| --- | --- | --- | --- |
| **Data collection** |  |  |  |
| EM equipment | Titan Krios | Titan Krios | Titan Krios |
| Detector | K3 camera | K3 camera | K3 camera |
| Magnification | 22,500x | 22,500x | 22,500x |
| Voltage (kV) | 300 | 300 | 300 |
| Electron exposure ((e-/Å<sup>2</sup>)) | 60 | 50 | 60 |
| Defocus range(μm) | -0.8 to -2.5 | -0.8 to -2.5 | -0.8 to -2.5 |
| Pixel size(Å) | 0.53 | 0.53 | 0.53 |
| Symmetry imposed | Helical | D2 | D2 |
| Number of collected movies | 5658 | 6695 | 5658 |
| Initial particle images (no.) | 1229156 | 3629981 | 2717149 |
| Final particle images (no.) | 512330 | 481216 | 337634 |
| Map resolution (Å) | 3.0 | 2.8 | 2.7 |
| FSC threshold | 0.143 | 0.143 | 0.143 |
| Map resolution range (Å) | 2.6 - 8.1 | 2.4 - 9.1 | 2.4 - 8.7 |
| **Refinement** |  |  |  |
| Initial model used (PDB code) | 2J5T | 2J5T | 2J5T |
| Model resolution (Å) | 3.1 | 2.8 | 2.8 |
| FSC threshold | 0.5 | 0.5 | 0.5 |
| Map sharpening B-factor(Å<sup>2</sup>) | -80 | -90 | -85 |
| Model composition |  |  |  |
| Non-hydrogen atoms | 21408 | 22070 | 22273 |
| Protein residues | 2848 | 2848 | 2848 |
| Ligands | 0 | Proline:8 | 0 |
| Waters | 0 | 0 | 865 |
| Ions | 0 | 0 | 0 |
| B factors(Å<sup>2</sup>) |  |  |  |
| Protein | 137.6 | 127.5 | 112.5 |
| Ligand | - | 126.2 | - |
| Water | - | 124.1 | 108.9 |
| R.m.s. deviations |  |  |  |
| Bond lengths (Å) | 0.003 | 0.003 | 0.002 |
| Bond angles (°) | 0.534 | 0.459 | 0.489 |
| Validation |  |  |  |
| MolProbity score | 1.38 | 1.62 | 1.73 |
| Clashscore | 5.82 | 5.48 | 7.69 |
| Poor rotamers (%) | 1.22 | 2.65 | 2.53 |
| Ramachandran plot |  |  |  |
| Favored (%) | 99.33 | 99.04 | 99.08 |
| Allowed (%) | 0.67 | 0.96 | 0.92 |
| Disallowed (%) | 0 | 0 | 0 |

**Supplementary Figure 1:**
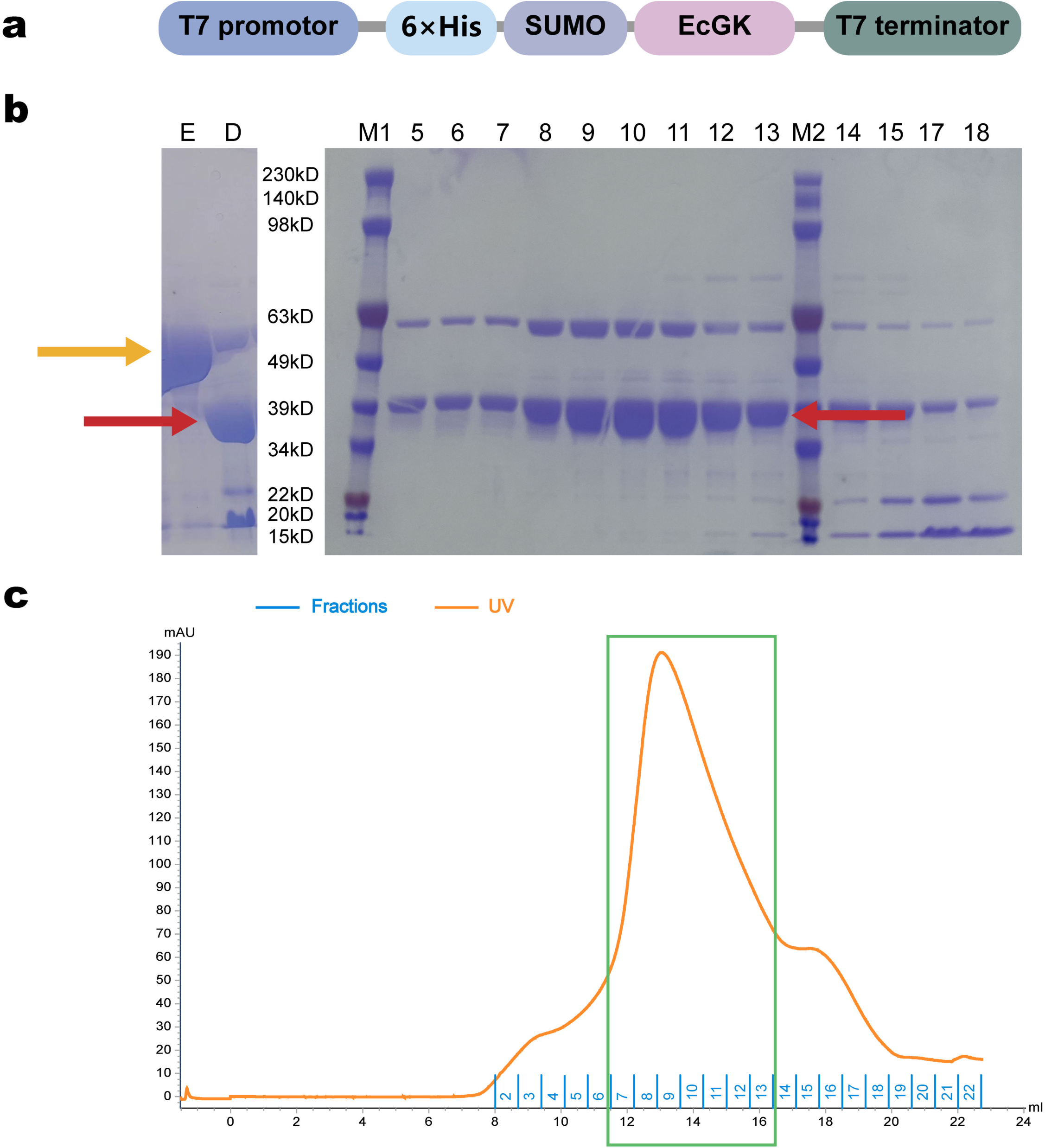
Qualification of EcGK during one purification experiment. a. Gene construction for EcGK expression vector. EcGK was fused with N-terminal His_6_-SUMO tag. Expression was driven by T7 promotor. b. SDS-PAGE of ecGK at different purification stages. The band corresponding to SUMO-EcGK fusion protein (about 51.49 kDa) is indicated with yellow arrow. The bands corresponding to EcGK protein (about 39.06 kDa) are indicated with red arrows. Different fractions during purification are represented with characters on the top of panel b. “E”: Elution from Ni^2+^-NTA beads. “D”: Elution after ULP1 digestion. “M1” and “M2”: Standard protein markers. The corresponding molecular weights are labeled at the sides of the bands. “5” to “18”: Different fractions after purification with size exclusion chromatography (Superose™ 6 Increase 10/300 GL). c. Elution profile from size exclusion chromatography. The orange curve indicates UV 280 nm absorbance. The fractions are numbered in blue above X-axis. The fraction numbers are the same as those in panel b. in this experiment we collected fraction 7-13 (indicated as green box) as final product.

**Supplementary Figure 2.**
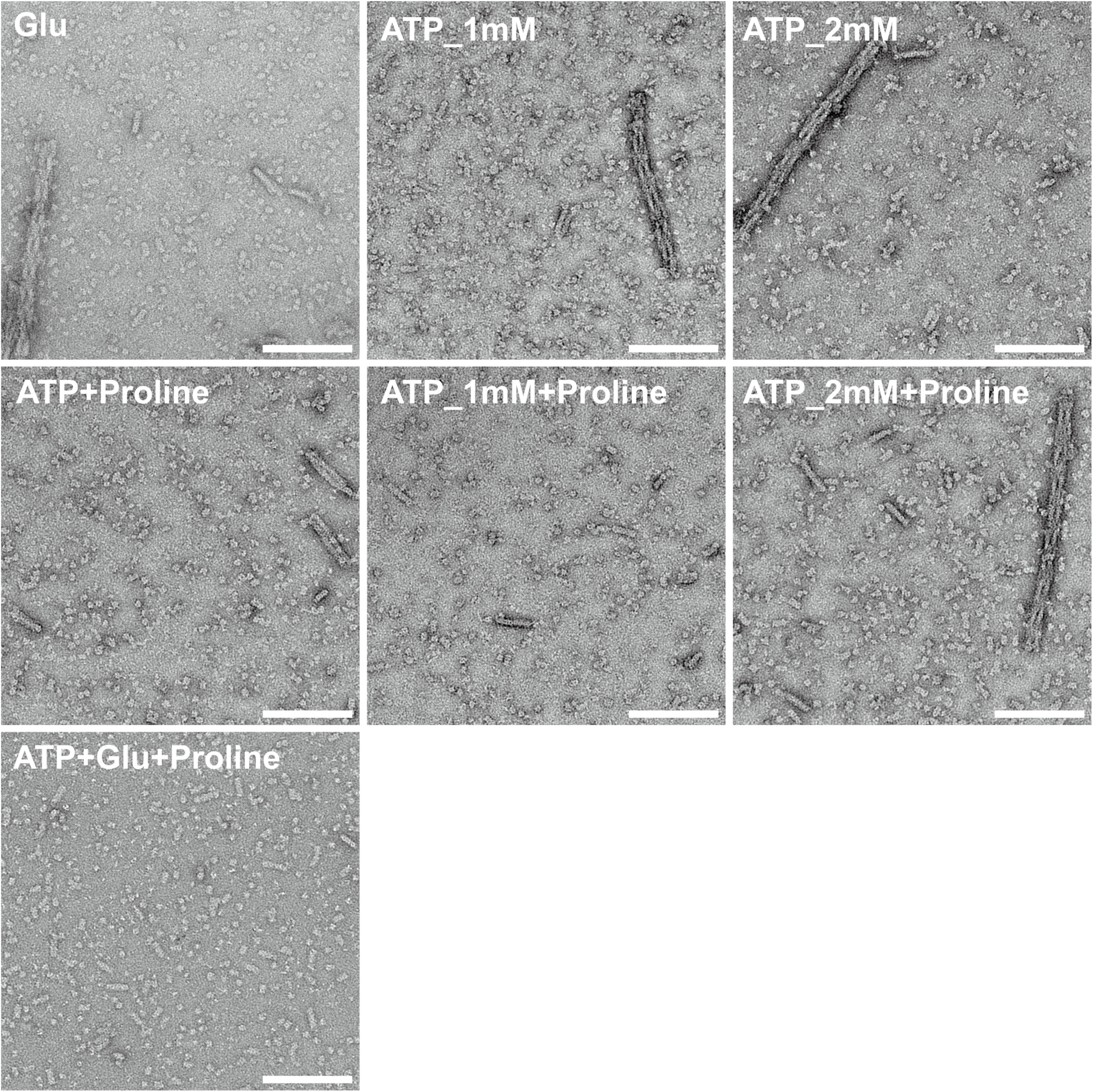
Negative staining results of EcGK at different conditions. Negative staining results of EcGK at different conditions. The concentration used: proline 0.5mM, glutamate 20mM, ATP 5mM. Scale bar: 150 nm.

**Supplementary Figure 3.**
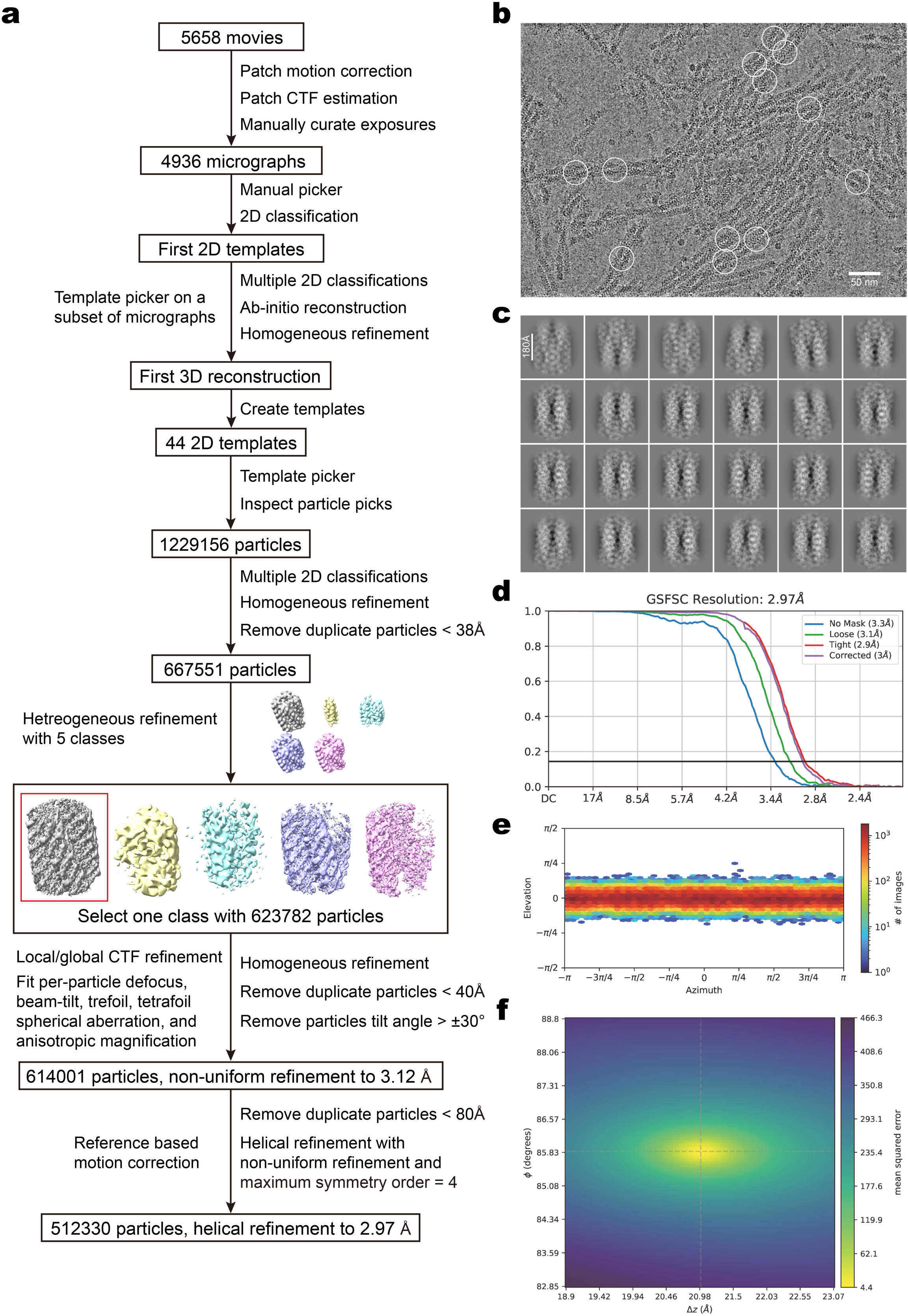
Reconstruction of apo-bundle structure. a. Data processing workflow. b. Representative micrograph after motion correction. Some apo-bundle particles are labeled with white circles. c. Representative 2D classes average from final 2D classification. d. Fourier shell correlation (FSC) curves for final 3D reconstruction. e. Particles angular distribution of final 3D reconstruction. f. Helical symmetry search results of final 3D reorganization.

**Supplementary Figure 4.**
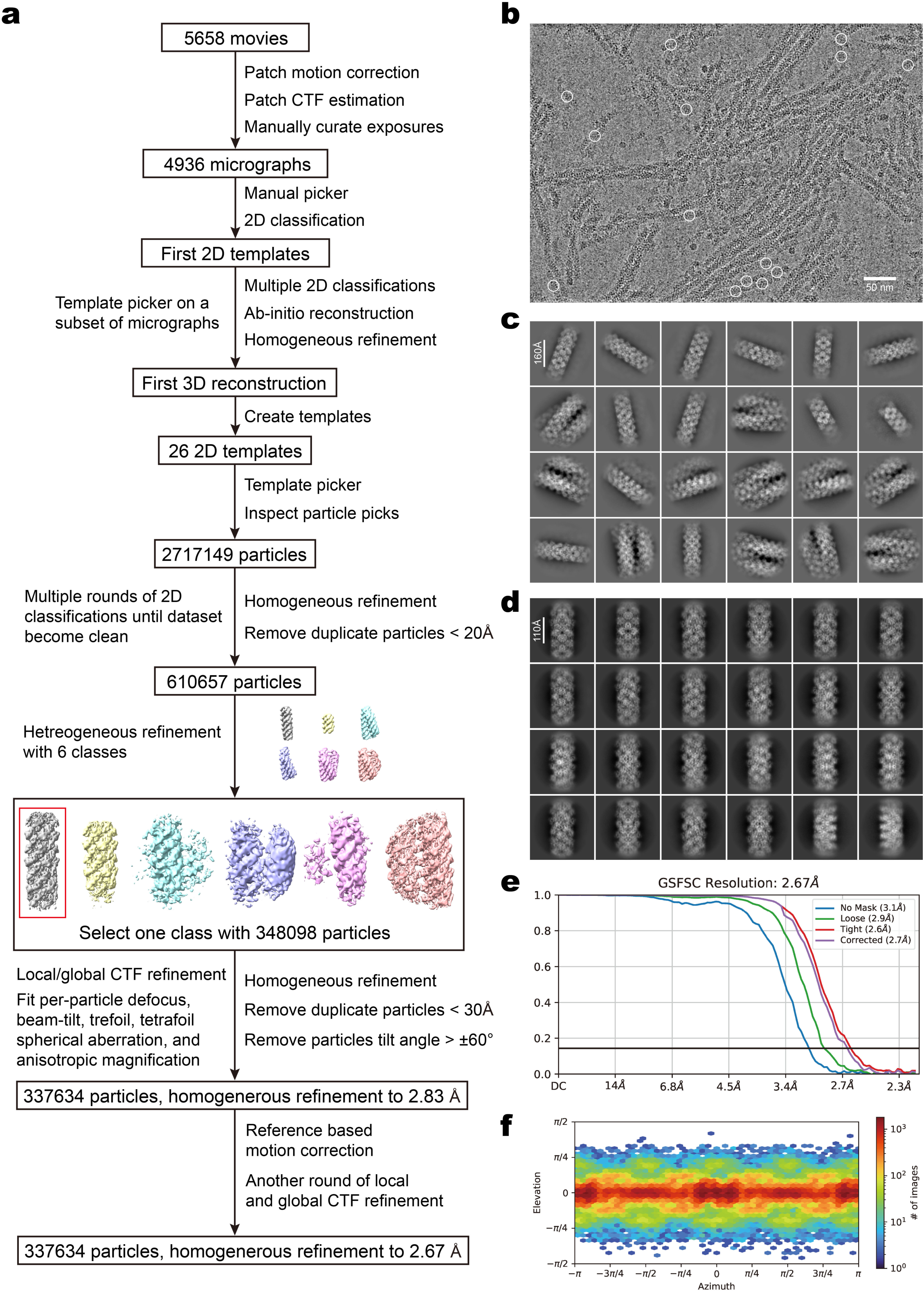
Reconstruction of apo-filament structure. a. Data processing workflow. b. Representative micrograph after motion correction. This is the same as Figure S3 b. Some apo-filament particles are labeled with white circles. c. Representative 2D classes average at beginning. d. Representative 2D classes average from final 2D classification. e. Fourier shell correlation (FSC) curves for final 3D reconstruction. f. Particles angular distribution of final 3D reconstruction.

**Supplementary Figure 5.**
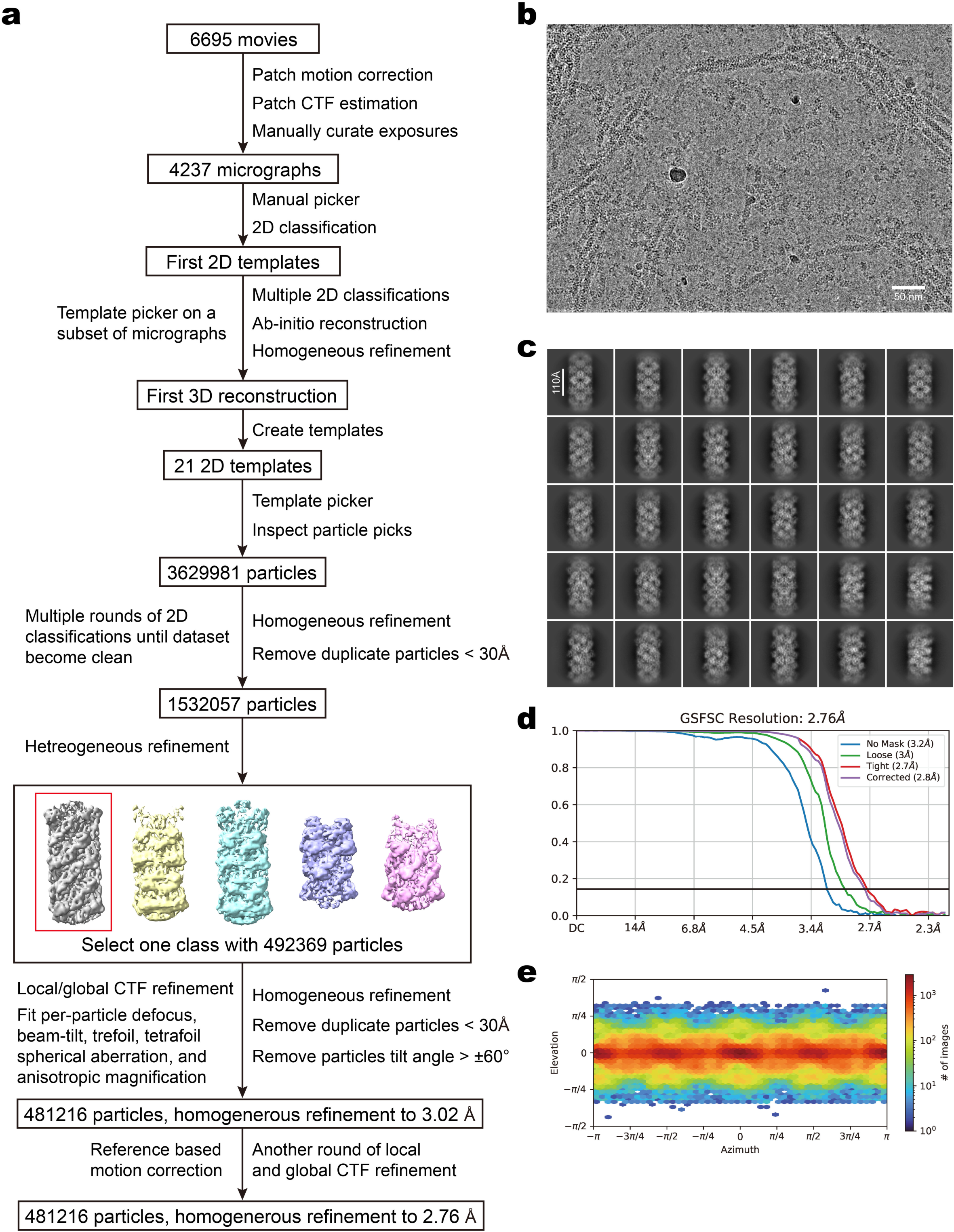
Reconstruction of proline-filament structure. a. Data processing workflow. b. Representative micrograph after motion correction. c. Representative 2D classes average from final 2D classification. d. Fourier shell correlation (FSC) curves for final 3D reconstruction. e. Particles angular distribution of final 3D reconstruction.

**Supplementary Figure 6.**
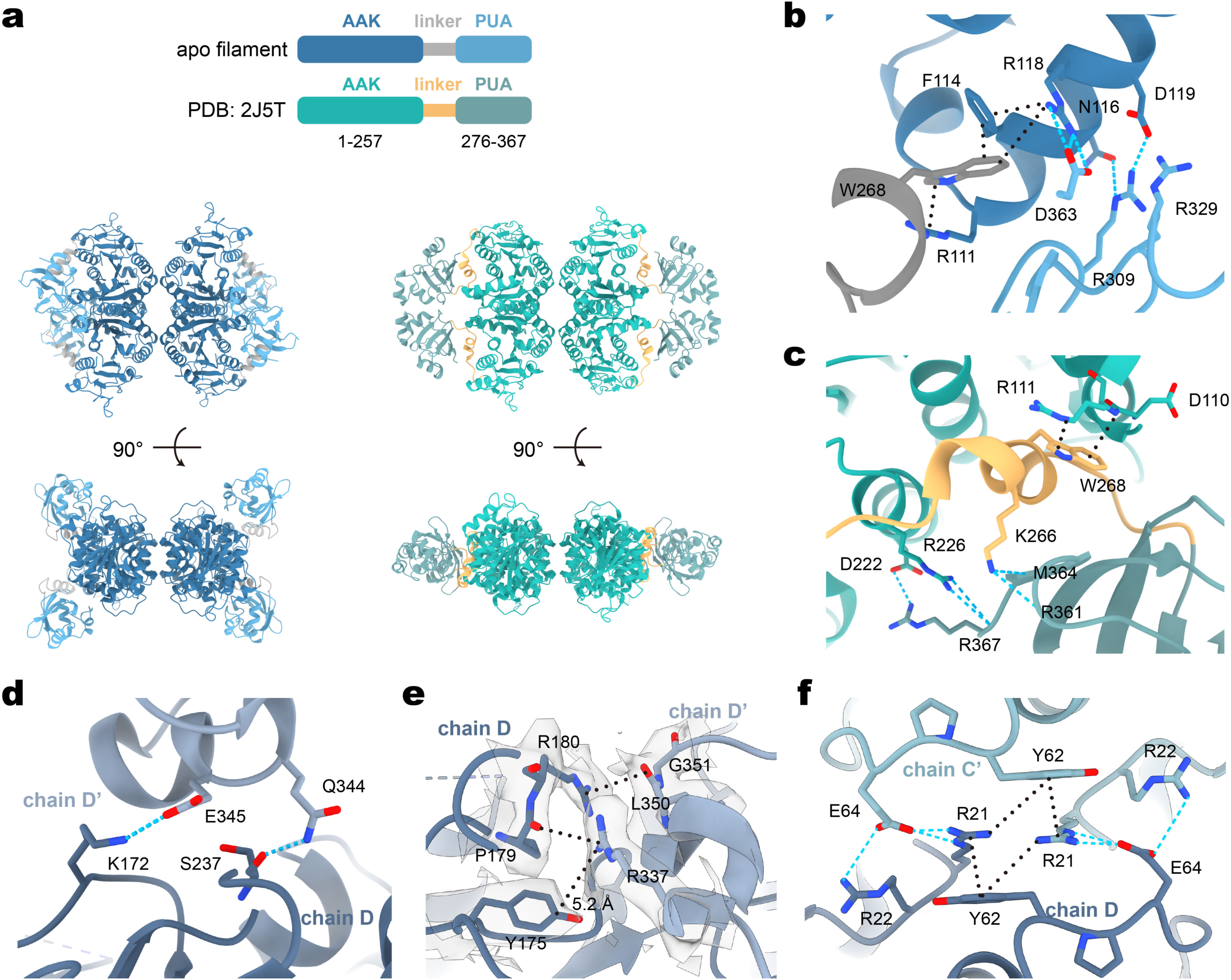
Detailed structural analysis of EcGK apo-filament. a. Comparison of tetramer from EcGK apo-filament and published model 2J5T. b-c: Detailed interactions analysis within b: apo-filament monomer; c: 2J5T monomer. AAK domain, linker, and PUA domain are color as the same scheme in panel a. d-f: Detailed interactions analysis at helical interfaces. Hydrogen bonding/electrostatic interactions are indicated by blue dashed lines. π-π/cation-π interactions are indicated by black dotted lines.

**Supplementary Figure 7.**
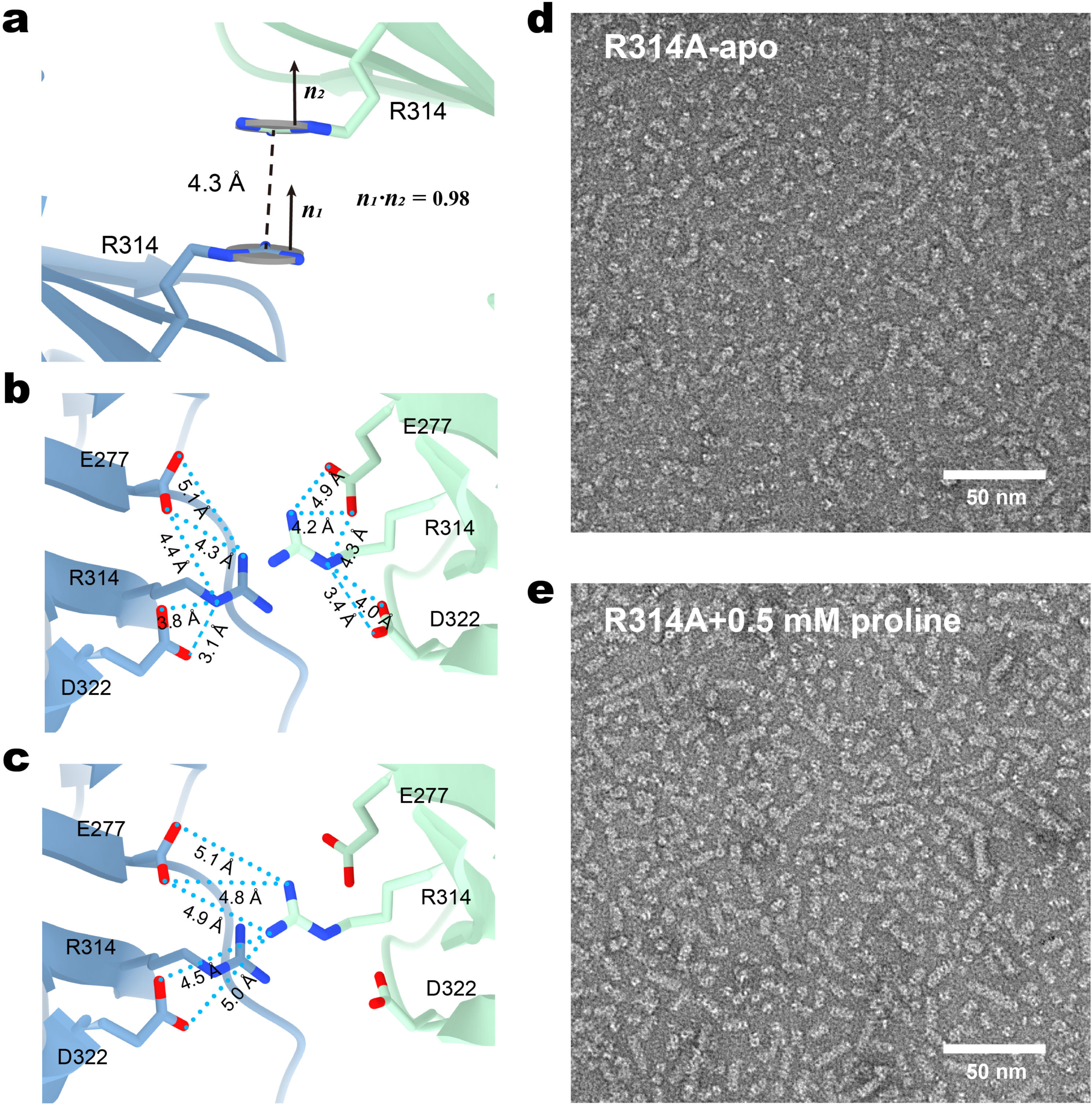
Analysis and validation of bundle interface. a. Geometric analysis of R314 on bundle interface. The planes of R314 guanidine groups are fitted with grey discs. Distance between two CZ atoms are measured. Normal vectors for the two planes are labeled as ***n_1_*** and ***n_2_***. b-c: Detailed interactions analysis at bundle interface. Possible electrostatic interactions are indicated by blue dashed lines. d-e: Negative staining results of EcGK-R314A mutant at different conditions.

**Supplementary Figure 8.**
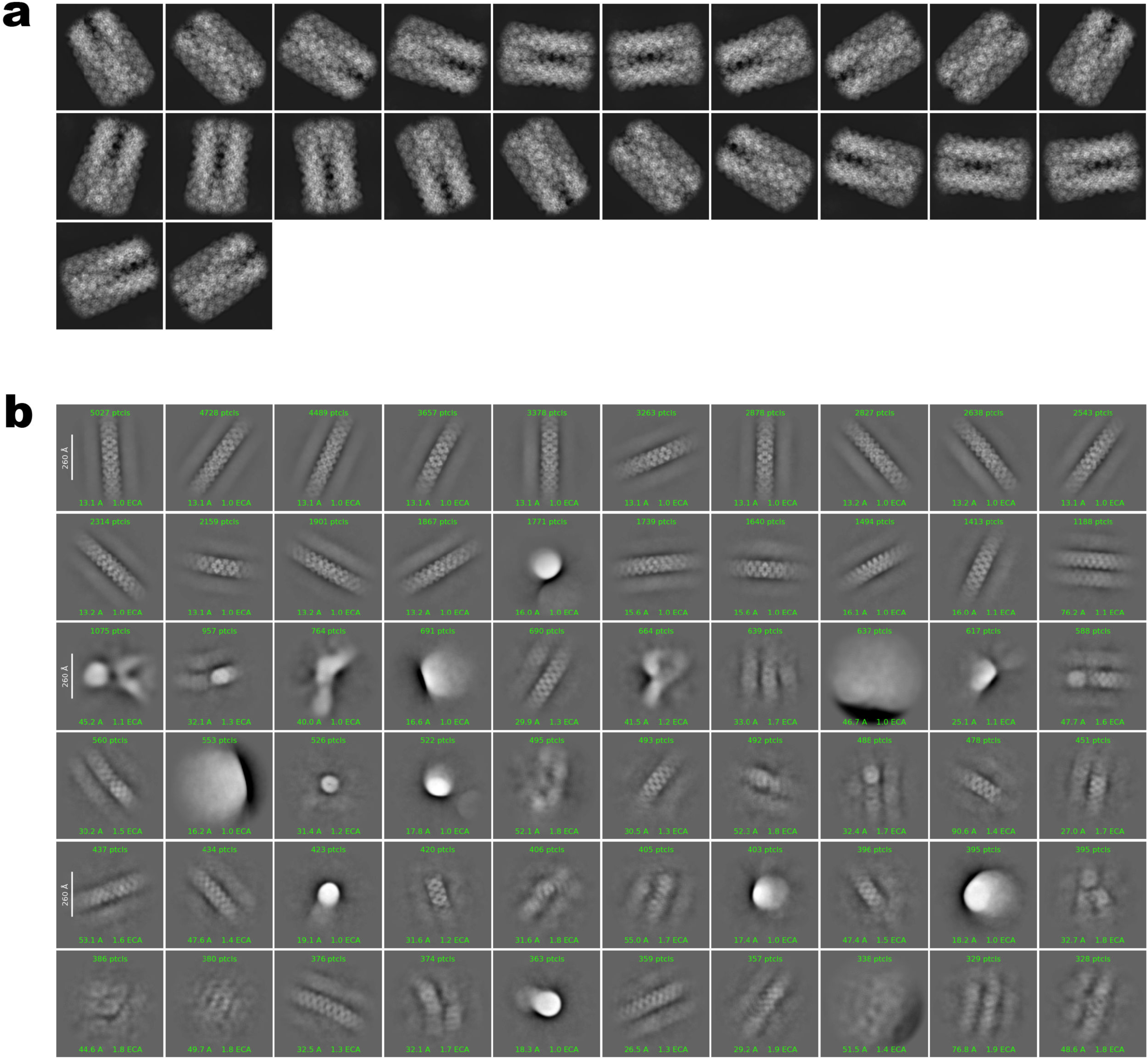
Attempt on picking bundles in proline sample. a. 2D templates used for picking bundles. b. 2D classification results of picked particles.

**Supplementary Figure 9.**
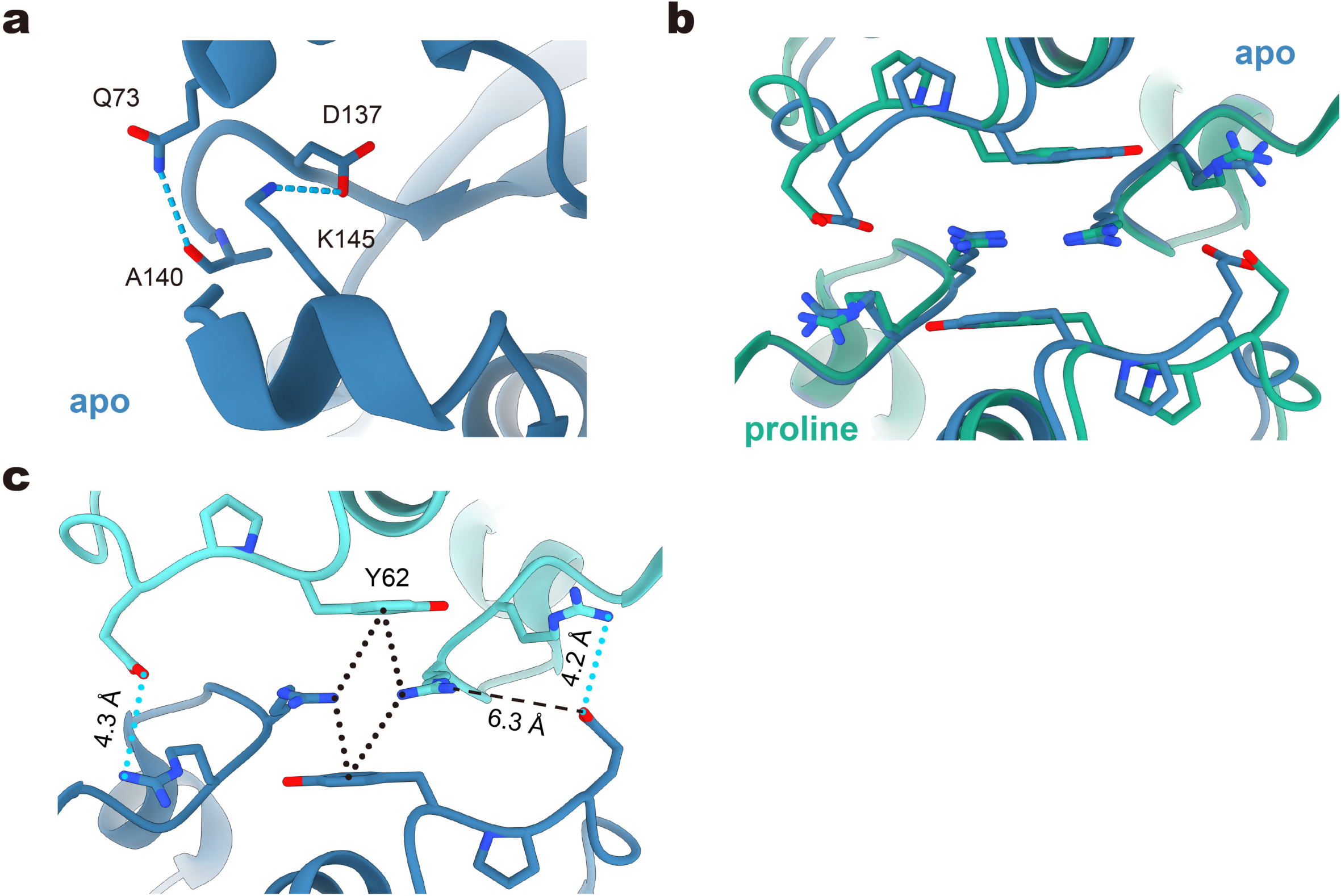
Detailed structural analysis of EcGK proline-filament interface. a-b: Detailed interactions analysis at proline-filament helical interfaces. c. Comparison of filament interface II between apo-filament and proline-filament.

**Supplementary Figure 10.**
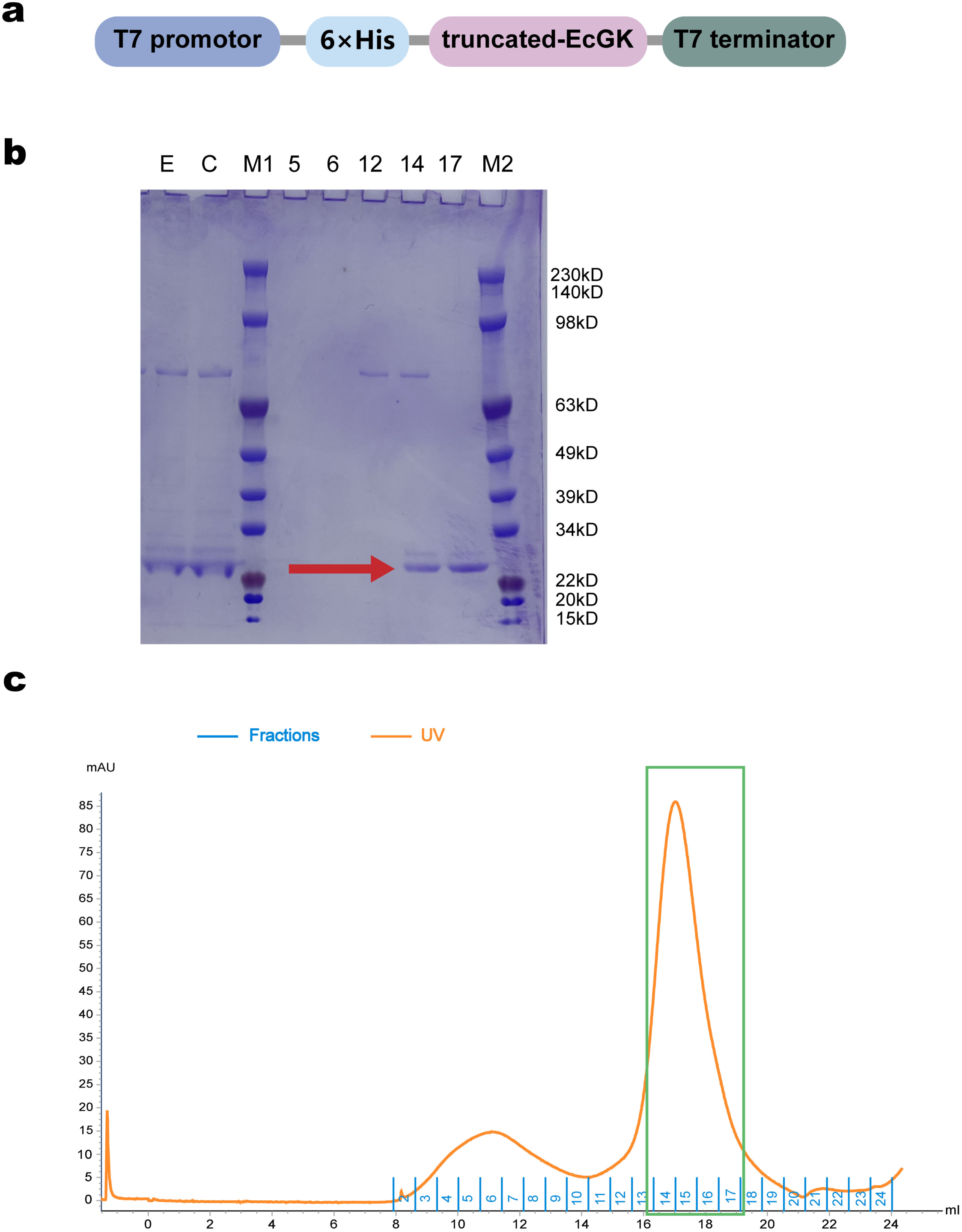
Qualification of truncated-EcGK during one purification experiment. a. Gene construction for truncated-EcGK expression vector. Truncated-EcGK was fused with N-terminal His6 tag. Expression was driven by T7 promotor. b. SDS-PAGE of truncated-EcGK at different purification stages. The band corresponding to truncated-EcGK fusion protein (about 30.22kDa) is indicated with red arrow. Different fractions during purification are represented with characters on the top of panel b. “E”: Elution from Ni2+-NTA beads. “C”: Elution after centrifugation. “M1” and “M2”: Standard protein markers. The corresponding molecular weights are labeled at the sides of the bands. “5” to “17”: Different fractions after purification with size exclusion chromatography (Superose™ 6 Increase 10/300 GL). c. Elution profile from size exclusion chromatography. The orange curve indicates UV 280 nm absorbance. The fractions are numbered in blue above X-axis. The fraction numbers are the same as those in panel b. In this experiment we collected fraction 14-17 (indicated as green box) as final product.

## Notes

### Competing Interest Statement

The authors have declared no competing interest.

